# Biological and clinical insights from genetics of insomnia symptoms

**DOI:** 10.1101/257956

**Authors:** Jacqueline M Lane, Samuel Jones, Hassan S Dashti, Andrew R Wood, Krishna Aragam, Vincent T. van Hees, Ben Brumpton, Bendik Winsvold, Heming Wang, Jack Bowden, Yanwei Song, Krunal Patel, Simon G Anderson, Robin Beaumont, David A Bechtold, Brian Cade, Sek Kathiresan, Max A Little, Annemarie I Luik, Andrew S Loudon, Shaun Purcell, Rebecca C Richmond, Frank AJL Scheer, Jessica Tyrrell, John Winkelman, HUNT All In Sleep, Linn B Strand, Jonas B. Nielsen, Cristen J. Willer, Susan Redline, Kai Spiegelhalder, Simon D Kyle, David W Ray, John-Anker Zwart, Kristian Hveem, Timothy M Frayling, Deborah Lawlor, Martin K Rutter, Michael N Weedon, Richa Saxena

**Affiliations:** Center for Genomic Medicine, Massachusetts General Hospital, Boston, MA, USA; Department of Anesthesia, Critical Care and Pain Medicine, Massachusetts General Hospital and Harvard Medical School, Boston, MA, USA; Broad Institute, Cambridge, MA, USA; Genetics of Complex Traits, University of Exeter Medical School, Exeter, United Kingdom; Cardiology Division, Massachusetts General Hospital and Harvard Medical School, Boston, MA, USA; Netherlands eScience Center, Amsterdam, The Netherlands; K.G. Jebsen Centre for Genetic Epidemiology, Department of Public Health and Nursing, Norwegian University of Science and Technology, Norway; MRC Integrative Epidemiology Unit at the University of Bristol, UK; Department of Thoracic and Occupational Medicine, St. Olavs Hospital, Trondheim University Hospital, Trondheim, Norway; Division of Sleep and Circadian Disorders, Department of Medicine, Brigham and Women's Hospital; Division of Sleep Medicine, Harvard Medical School, University of Bristol, UK; Population Health Sciences, Bristol Medical School, University of Bristol, UK; Northeastern University College of Science, 206 Mugar Life Sciences, 360 Huntington Avenue, Boston, MA 02115; Division of Cardiovascular Sciences, School of Medical Sciences, Faculty of Biology, Medicine and Health, The University of Manchester, Manchester, UK (SA); The George Institute for Global Health, University of Oxford, Oxford Martin School, 34 Broad Street Oxford OX1 3BD, UK (SA); Faculty of Biology, Medicine and Health, University of Manchester, Manchester, UK; Division of Endocrinology, Diabetes & Gastroenterology, School of Medical Sciences, Faculty of Biology, Medicine and Health, University of Manchester, UK; Cardiovascular Research Center, Massachusetts General Hospital, 185 Cambridge Street, Boston, Massachusetts 02114, USA; Department of Mathematics, Aston University, Birmingham, UK; Media Lab, Massachusetts Institute of Technology, Cambridge, Massachusetts, USA; Sleep and Circadian Neuroscience Institute, Nuffield Department of Clinical Neurosciences, University of Oxford, Oxford, UK; Department of Psychiatry, Brigham & Women's Hospital, Harvard Medical School, Boston, MA, 02115; School of Social and Community Medicine, University of Bristol, Bristol, UK; Medical Research Council Integrative Epidemiology Unit, University of Bristol, Oakfield House, Bristol, BS8 2BN UK; Departments of Psychiatry and Neurology, Massachusetts General Hospital, Boston, MA, USA; FORMI and Department of Neurology, Oslo University Hospital, Oslo, Norway; Division of Cardiovascular Medicine, Department of Internal Medicine, University of Michigan, Ann Arbor, Michigan, USA; Department of Human Genetics, University of Michigan, Ann Arbor, Michigan, USA; Department of Computational Medicine and Bioinformatics, University of Michigan, Ann Arbor, Michigan, USA; Departments of Medicine, Brigham and Women's Hospital and Beth Israel Deaconess Medical Center, Harvard Medical School, Boston; Clinic for Psychiatry and Psychotherapy, Medical Centre - University of Freiburg, Faculty of Medicine, University of Freiburg, Germany; Manchester Diabetes Centre, Manchester University NHS Foundation Trust, Manchester Academic Health Science Centre, Manchester, UK

## Abstract

Insomnia is a common disorder linked with adverse long-term medical and psychiatric outcomes, but underlying pathophysiological processes and causal relationships with disease are poorly understood. Here we identify 57 loci for self-reported insomnia symptoms in the UK Biobank (n=453,379) and confirm their impact on self-reported insomnia symptoms in the HUNT study (n=14,923 cases, 47,610 controls), physician diagnosed insomnia in Partners Biobank (n=2,217 cases, 14,240 controls), and accelerometer-derived measures of sleep efficiency and sleep duration in the UK Biobank (n=83,726). Our results suggest enrichment of genes involved in ubiquitin-mediated proteolysis, phototransduction and muscle development pathways and of genes expressed in multiple brain regions, skeletal muscle and adrenal gland. Evidence of shared genetic factors is found between frequent insomnia symptoms and restless legs syndrome, aging, cardio-metabolic, behavioral, psychiatric and reproductive traits. Evidence is found for a possible causal link between insomnia symptoms and coronary heart disease, depressive symptoms and subjective well-being.

**One Sentence Summary:** We identify 57 genomic regions associated with insomnia pointing to the involvement of phototransduction and ubiquitination and potential causal links to CAD and depression.

---

Insomnia disorder, defined by persistent difficulty in initiating or maintaining sleep, and corresponding daytime dysfunction, occurs in roughly 10-20% of the population, and leads to high socioeconomic costs and substantial lifetime morbidity^1^. Up to one-third of the population experience transient insomnia symptoms at any given time^2^. Longitudinal studies suggest that insomnia increases the risk for developing anxiety disorders, alcohol abuse, major depression and cardio-metabolic disease^3^. Despite its high prevalence and a hypothesized strong bidirectional link between insomnia and psychiatric disorders, little is known about underlying pathophysiologic mechanisms. Cognitive-behavioral therapies are the recommended first-line treatment approach but access is limited^4,5^. Common drug treatments target synaptic neurotransmission (via GABAergic pathways), cortical arousal (via histamine receptors), or the melatonin system, but these drugs have variable effectiveness, may be habit forming and have important side effects^6,7^. A better understanding of the etiology and pathophysiological processes would enable identification of new personalized therapeutic strategies for insomnia. Family-based heritability estimates suggest that insomnia has a genetic component (22%–25%). Recent GWAS in the first release of genetic data from the UK Biobank have reported four loci for insomnia symptoms (at *MEIS1, TMEM132E, CYCL1* and SCFD2)^9,10^, but insights into underlying biological pathways and causal genetic links with disease are limited.

In this study, we aimed to 1) discover novel genetic loci for self-reported insomnia symptoms using GWAS, 2) validate findings in independent clinical and population samples and in participants with activity monitor derived measures of sleep patterns and, 3) gain biological insights by gene, pathway and tissue-enrichment analyses and bioinformatic annotation, 4) investigate shared genetics with behavioral and disease traits, and 5) test for causal links between insomnia and relevant disease/traits.

In UK Biobank participants of European ancestry (n=453,379), 29% self-reported frequent insomnia symptoms, with a higher prevalence in women than men (32% vs. 24%). Consistent with previous studies, insomnia symptoms were more prevalent in older participants, shift workers and those with shorter self-reported sleep duration (Supplementary Table 1).

We performed two parallel GWAS in participants self-reporting insomnia symptoms 1) frequent insomnia symptoms: never/rarely vs. usually insomnia symptoms, n=129,270 2) any insomnia symptoms: never/rarely vs. sometimes/usually insomnia symptoms, n=345,022 cases using 14,661,600 genetic variants across the autosomes and X chromosome. We identified 57 association signals (Fig. 1, Supplementary Table 2, Supplementary Fig.1-2). Of these, 20 loci were identified in both analyses, 28 loci were identified only in analysis of frequent insomnia symptoms, and 9 only in analysis of any insomnia symptoms (Supplementary Table 2). Conditional analyses identified no secondary association signals. The 57 genetic associations were independent of established or putative insomnia risk factors, as sensitivity analyses adjusting for BMI, lifestyle, caffeine consumption, and depression or recent stress did not notably alter the magnitude or direction of effect estimates (Supplementary Table 3). The *MEIS1* association signal identified in the interim release of the UK Biobank was confirmed in the remainder of the UK Biobank sample (excluding the interim subjects and relatives of interim subjects) (n=75,508 cases of frequent insomnia symptoms and 64,403 controls; rs113851554 T OR [95% CI] 1.19 [1.15-1.23, p=1.5 x10^-21^), and nominal replication was seen for the previously reported *CYCL1* signal (*p=9.0* x10^-3^) The *TMEM132E* and *SCFD2* signals showed concordant direction of effect with the initial subsample, but were not significant, perhaps reflecting selection bias in the initial subsample, with a high respiratory disease burden from the BILEVE study^11^. No other findings from previous candidate gene association studies or smaller GWAS were confirmed (Supplementary Table 4).

**Figure 1.**
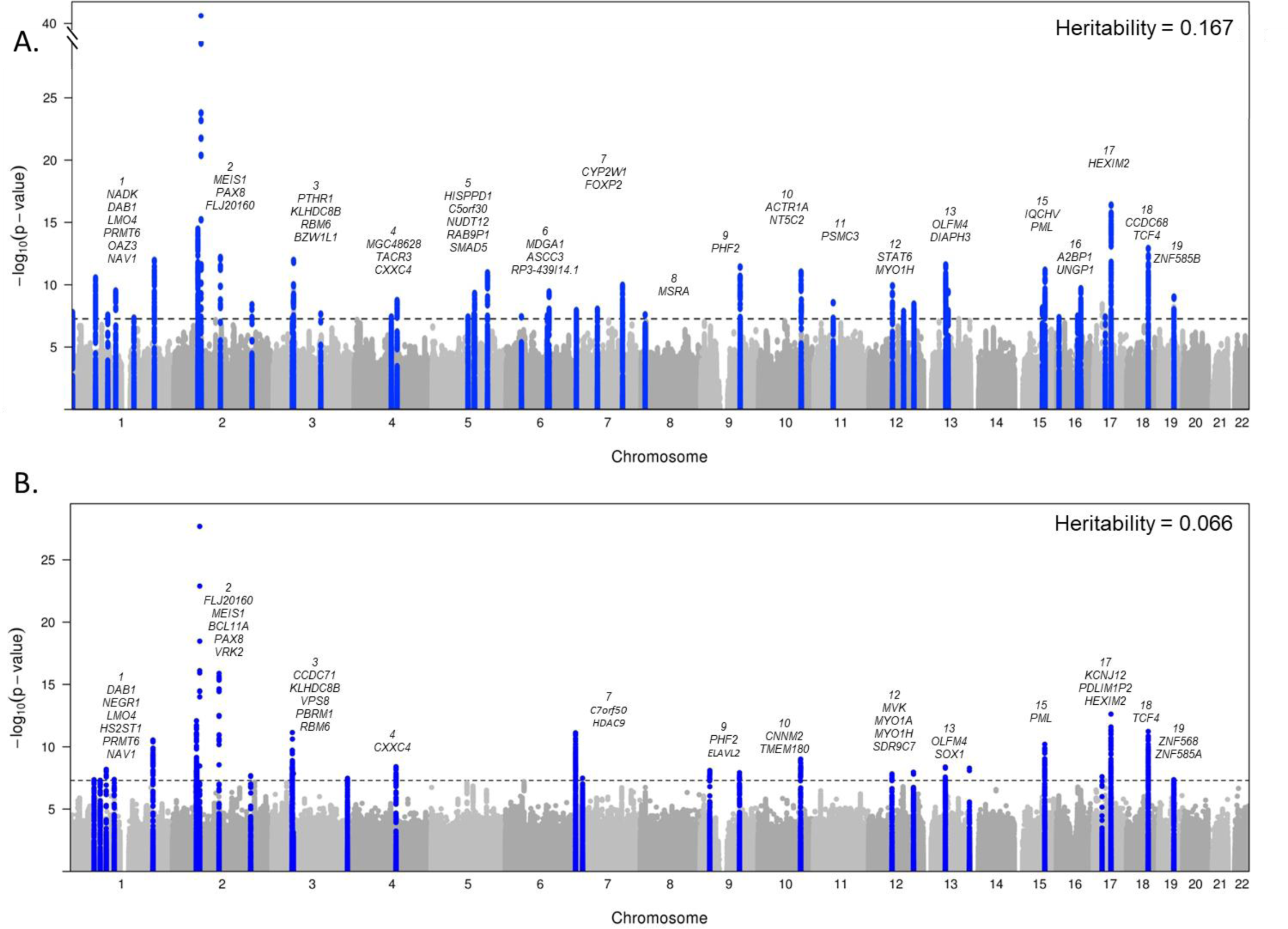
Manhattan plots for genome-wide association analysis of frequent (A) and any (B) insomnia symptoms. Dotted line is genome-wide significant (5×10^-8^). Heritability estimates were calculated using BOLT-REML. Chromosomes are annotated with the nearest gene to each association signal.

Secondary GWAS excluding current shift workers or individuals reporting hypnotic, anti-anxiolytic or psychiatric medication usage, and/or with selected chronic diseases or psychiatric illnesses, (excluding n= 76,470 participants) revealed strong pair-wise genetic correlation to the primary GWAS (r_g_~1) and did not identify any additional association signals (Supplementary Fig. 1-3). Thus, biological processes underlying pathophysiology of insomnia symptoms may be common between the general population and those with co-morbidities, in accordance with the recent clinical reclassification of primary and secondary insomnia diagnoses into an insomnia disorder^12^.

The prevalence of insomnia symptoms varies by sex, therefore we performed secondary sex stratified GWAS for both frequent and any insomnia symptoms. Thirteen loci were found (8 in women and 5 in men), of which seven demonstrated evidence of sex interactions (*p*_sex-int_<3x10^-4^; with stronger effects in women at *KRT8P18, NT5C2, NMT1, CCDC148, C11ORF49* and stronger effects in men at *CADM1* and *SLC8A3;* Supplementary Table 5). Effects in women were not modified by menopausal status (Supplementary Table 5). Furthermore, as described previously^9,10^, the genetic architecture for frequent insomnia symptoms differed by sex, with a genetic correlation between the stratified GWAS of r_g_=0.807 (Supplementary Fig.3).

Given the limitations of the self-report of insomnia symptoms^13^, we sought additional replication and validation of genetic association signals. First, we tested our 57 lead variants for association with self-reported insomnia symptoms in participants from the population-based HUNT study (n=14,923 cases, 47,610 controls; characteristics described in Supplementary Table 6)^14^. Replication was observed for the *MEIS1* variant, and 40/57 variants showed a consistent direction of effect across both studies (binomial test *p*=9 x 10^-4^)(Supplementary Table 7). A genetic risk score of 57 variants (GRS) weighted by effect estimates from the primary UK Biobank GWAS was also associated with insomnia symptoms in HUNT (OR [95%CI] 1.015 [1.01-1.02] per allele, p=2.71x10^-11^) (Table 1). Second, we tested for and found an association of the GRS with physician diagnosed insomnia in the Partners Biobank (n= 2,217 cases, 14,240 controls; OR [95%CI) 1.017 [1.007-1.027] per allele, 8.88x10^-4^; Table 1, Supplementary Table 8) Third, to investigate impact of genetic variants on objective sleep patterns, we tested the 57 lead variants for association with 8 activity-monitor measures of sleep fragmentation, duration and timing in a subset of the UK Biobank participants of European ancestry who had undergone 7 days of wrist-worn accelerometry (n=84,745, characteristics described in Supplementary Table 6). The lead *MEIS1* risk variant was associated with a higher number of sleep episodes, lower sleep efficiency, shorter sleep duration and later sleep timing (p<0.0008; Supplementary Table.9) The GRS was associated with reduced sleep efficiency (difference = -0.04 (0.01) % per allele; p=4 x 10^-14^), shorter sleep duration (difference = -0.25 (0.035) mins per allele; p=8 x 10^-13^) and greater day-to-day variability in sleep duration (difference = 0.077 (0.025) mins per allele; p=0.0017) but not with the number of sleep episodes or diurnal inactivity duration (Table 1, Supplementary Table 9).

**Table 1.**
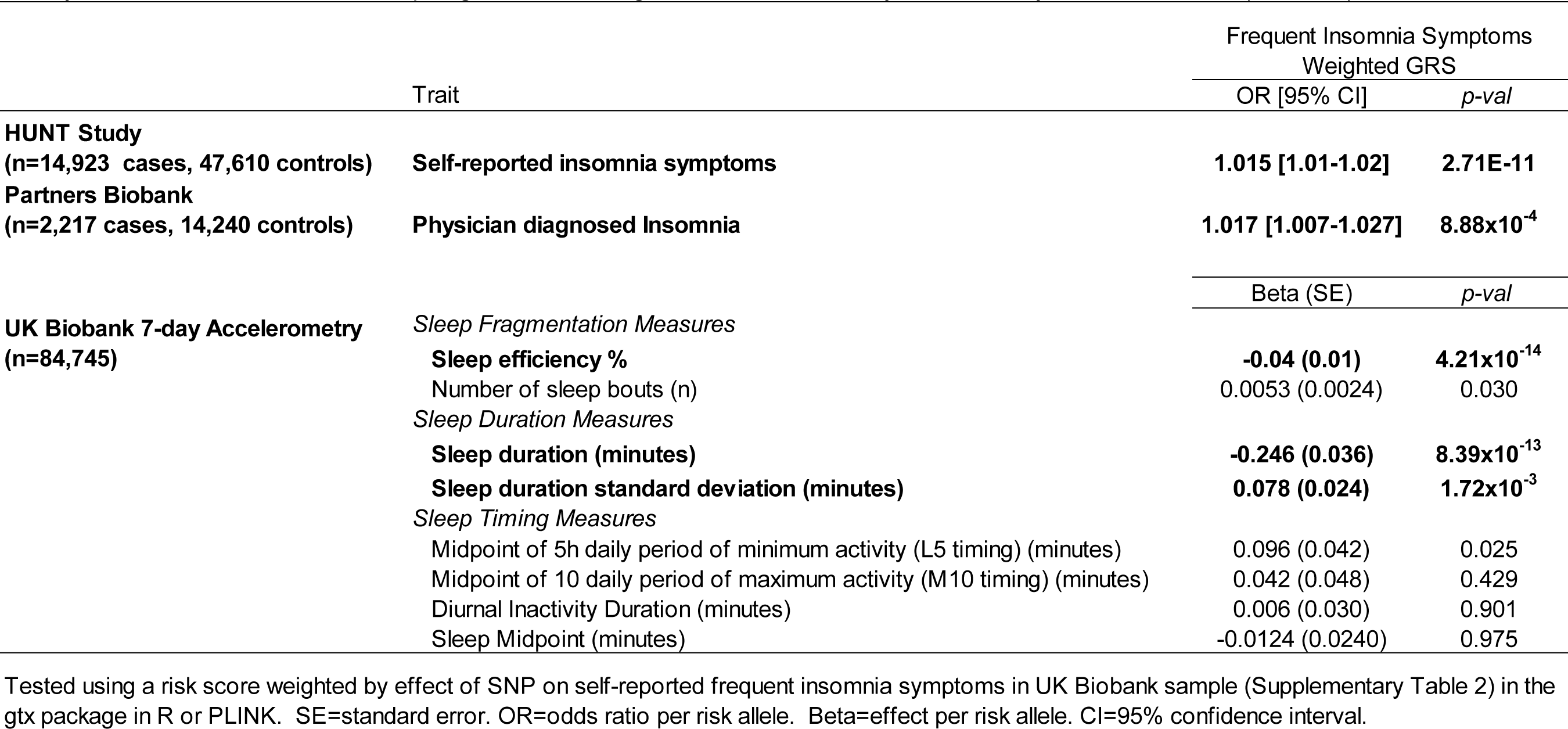
A risk score of genetic variants for self-reported insomnia symptoms (57 SNPs) associates with a) self-reported insomnia symptoms in the HUNT study (n=14,923 cases, 47,610 controls), b)physician diagnosed insomnia in the Partners Biobank (n=2,217 cases, 14,240 controls) and c) activity-monitor based measures of sleep fragmentation, timing and duration from 7 day accelerometry in the UK Biobank (n=84,745).

In order to gain insight into the probable causal variant underlying the 57 genetic association signals, we performed fine-mapping based on 1KG project linkage information using credible set analysis in PICS^15^ and identified 38 variants with a causal probability of 0.20 or greater (Supplementary Table 10). The majority of likely causal variants lie within introns (34%) or downstream of a gene (22%), consistent with previous literature demonstrating that non-coding variation causally influences the majority of phenotypic associations for complex traits (Supplementary Fig.4)^16^. This list includes missense variants located in NAD kinase *NADK* (N262K) and *MDGA1* (L61P) and rs324017, a SNP within the gene encoding transcriptional repressor *NAB1* (also known as EGR-1 binding protein) that is predicted to disrupt a binding site for EGR1 (Supplementary Table 10), a transcription factor involved in response to stress^17^ and in synaptic plasticity during REM sleep^17^ (Supplementary Table 9).

The 57 insomnia symptoms loci lie in genomic regions encompassing up to 236 genes, and a summary of annotations from public databases that link signals or genes within each locus to phenotypes in humans and model organisms is shown in Supplementary Table 11. Association signals at 14 loci directly overlapped with NHGRI GWAS signals (r^2^>0.7 in 1KG CEU) for one or more complex traits, with the insomnia symptoms risk allele associated with a greater risk of restless legs syndrome *(MEIS1, MAP2K5, TOX3)*, schizophrenia *(VRK2* region), Tourette’s syndrome or obsessive-compulsive disorder *(FLJ30838* region), and associated with higher systolic blood pressure *(NADK)*, greater carotid plaque burden (OAZ3), lower age at menarche (*LIN28B*) and influencing adiposity traits, height and educational attainment. Mutations in several genes under association peaks have been implicated in Mendelian cancers, connective tissue disorders, disorders with ocular and other sensory dysfunction and hypomagnesia, among others. Notably, experimental studies in mouse or fly models have implicated five genes at 3 associated loci in sleep regulation *(DVL1, LRP1, NR1H3, PRKAR2A*, and *SEMA3F).* Genes within 16 loci are known drug targets.

Two lead SNPs were associated with one or more of 3,144 human brain structure and function traits assessed in the UK Biobank (p<2.8x10^-7^, n=9,707; Oxford Brain Imaging Genetics Server)^18^, including rs1544637 (within a transcript *LOC642659)* with several large white matter tracts and with cingulate gyrus morphometry, which has previously been connected to insomnia^19^, and rs62158170 (near *PAX8)* with resting-state fMRI networks (Supplementary Fig.5).

Gene-based tests^20^ that aggregate genetic effects within a gene identified 135 genes associated with frequent insomnia symptoms (p≤2.29x10^-6^; Supplementary Table 12). These genes are enriched for expression in brain regions^9^ (Supplementary Table 13), including the cerebellum (p=1.3 x 10^-6^), frontal cortex (p= 1.3 x 10^-5^), anterior cingulated cortex (p=1.7 x 10^-5^), hypothalamus (p=2.2 x 10^-5^), basal ganglia (p=7.0 x 10^-4^), amygdala (p=3.4 x 10^-4^), and hippocampus (p=8.4 x 10^-4^), in line with previous reports in the literature linking these brain regions to insomnia^21,22^. Integration of gene expression data with GWAS using transcriptome-wide association analyses^23^ identified 24 genes for which insomnia-SNPs influence gene expression in one or more of 14 tissue types tested, including eight brain regions, muscles, peripheral nerves, whole blood, pituitary, thyroid and adrenal gland tissue (Supplementary Table 16).

SNP-based heritability of frequent insomnia symptoms was estimated to be h^2^ =16.7%^24^. Partitioning of heritability across tissue types^25,26^ showed enrichment in the central nervous system, adrenal/pancreas tissue lineages and skeletal muscle (*p*<10^-5^) (Supplementary Table 17). Additionally, partitioning across functional class implicated activation and repression of enhancers in the etiology of insomnia symptoms (Supplementary Table 17). Pathway and ontology analyses^20,27,28^ reveal a significant role for ubiquitin mediated proteolysis (p_bonf_=0.04; Supplementary Table 14), and suggestive roles for photo transduction and muscle tissue development and structures in the generation of frequent insomnia symptoms, with results consistent across several pathway and ontological databases (Supplementary Table 14, Supplementary Table 15). Evidence from model organisms supports the link between Cullin-3 mediated ubiquitination and sleep and circadian rhythms^29-32^. Furthermore, the restless legs syndrome gene *BTBD9* has been implicated as a substrate adaptor for the Cullin-3 class of E3 ubiquitin ligases^33^.

We investigated the genetic link between frequent insomnia symptoms and other behavioral and/or disease states. Based on previously identified genetic links between restless legs syndrome (RLS) and insomnia symptoms^9,10^, we tested a GRS of 20 SNPs for RLS^34^ and found association with frequent insomnia symptoms (OR=1.03 (1.02-1.04) per RLS risk allele,p=5.69x10^-53^; Supplementary Table 18). To test the proportion of variance frequent insomnia symptoms shares with other traits based on genetic overlap, we performed genome-wide genetic correlation analyses between our frequent insomnia symptoms GWAS and 233 traits with public GWAS summary statistics^25,26,35,36^ We found and strong positive genetic correlations (p≤2×10^-3^) between frequent insomnia symptoms and adiposity traits, coronary artery disease, neuroticism, smoking behavior and depressive symptoms and disorder. Strong negative genetic correlations were observed with self-reported sleep duration, subjective well-being, cognitive measures, proxy longevity measures and reproductive traits (Figure 2, Supplementary Table 19). These genetic links persisted with our frequent insomnia symptoms GWAS excluding subjects with chronic and psychiatric illnesses (Supplementary Table 19), indicating a relationship not driven by the presence of concomitant conditions.

**Figure 2.**
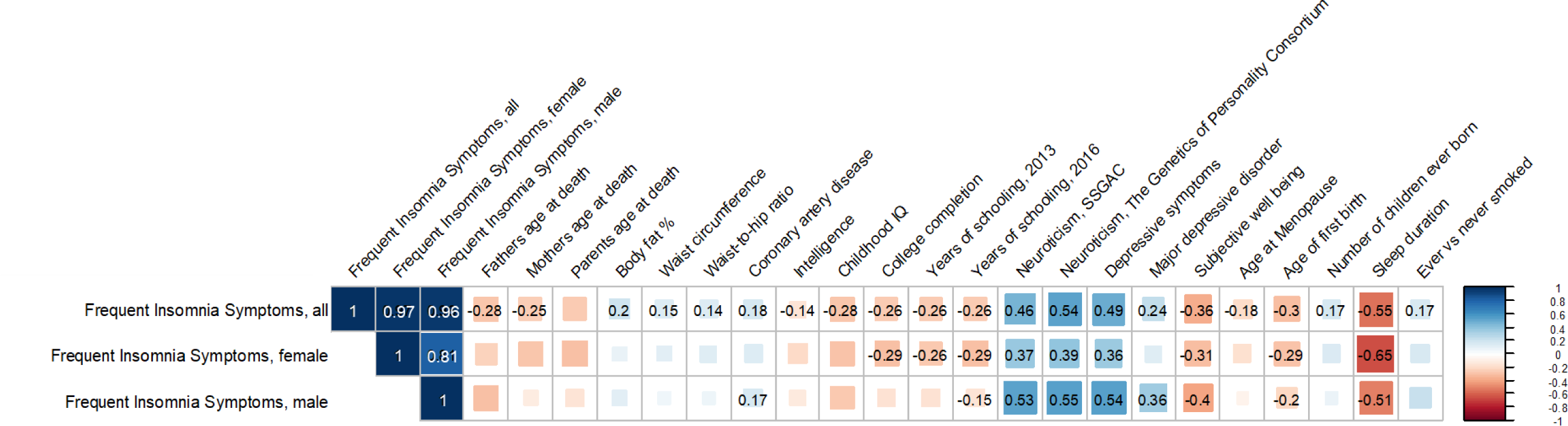
Shared genetic architecture between frequent insomnia symptoms and behavioral and disease traits. LD-score regression estimates of genetic correlation (r_G_) of frequent insomnia symptoms are compared with the summary statistics from 224 publicly available genome-wide association studies of psychiatric and metabolic disorders, immune diseases, and other traits of natural variation. Blue, positive genetic correlation; red, negative genetic correlation, r_G_ values displayed for significant correlations. Larger squares correspond to more significant p-values. Genetic correlations that are significantly different from zero after Bonferroni correction are shown on the plot, after Bonferroni correction p-value cut-off is 0.0002. All genetic correlations in this report can be found in tabular form in Supplementary Table 19. Abbreviations: IQ=intelligence quotient.

To test for causal links between frequent insomnia symptoms and seven clusters of genetically correlated traits, we performed Mendelian Randomization analyses mostly within a two-sample summary data framework leveraging the genotype disease association from publicly available sources to make inferences about causality. Using our main inverse variance (IVW) MR approach^37^ (Fig 3, Supplementary Table 20), we find evidence of a causal association (IVW p<0.001) between frequent insomnia symptoms and coronary artery disease (OR=2.15 (95% CI 1.38-3.35) for CAD risk with genetically instrumented case status for frequent insomnia symptoms using all 57 variants), for reduced subjective well-being with difference in mean SD units of -0.29 (s.e. 0.06) and for greater depressive symptoms with difference in mean SD units of 0.42 (s.e. 0.08), with a consistent direction and similar magnitude of effect in sensitivity analyses using MR-Egger^38^ and weighted median methods (Fig. 3, Supplementary Table 20)^39^. We find similar results using effect estimates from our GWAS excluding those with preexisting conditions. We validated the possible casual association between insomnia and CAD by performing a 1-sample MR in the UK Biobank (N= cases 23,980 and 361,706 controls, OR [95%CI] 2.95 [2.18 - 3.99], p=2.3 x 10^-12^)(Supplementary Table 20 and Supplementary Fig. 6).We find no evidence of reverse causality between CAD and insomnia. The causal association between insomnia status and coronary artery disease in UK Biobank is consistent with the robust empirical evidence seen in prospective studies and meta-analyses^3,40^.

**Figure 3.**
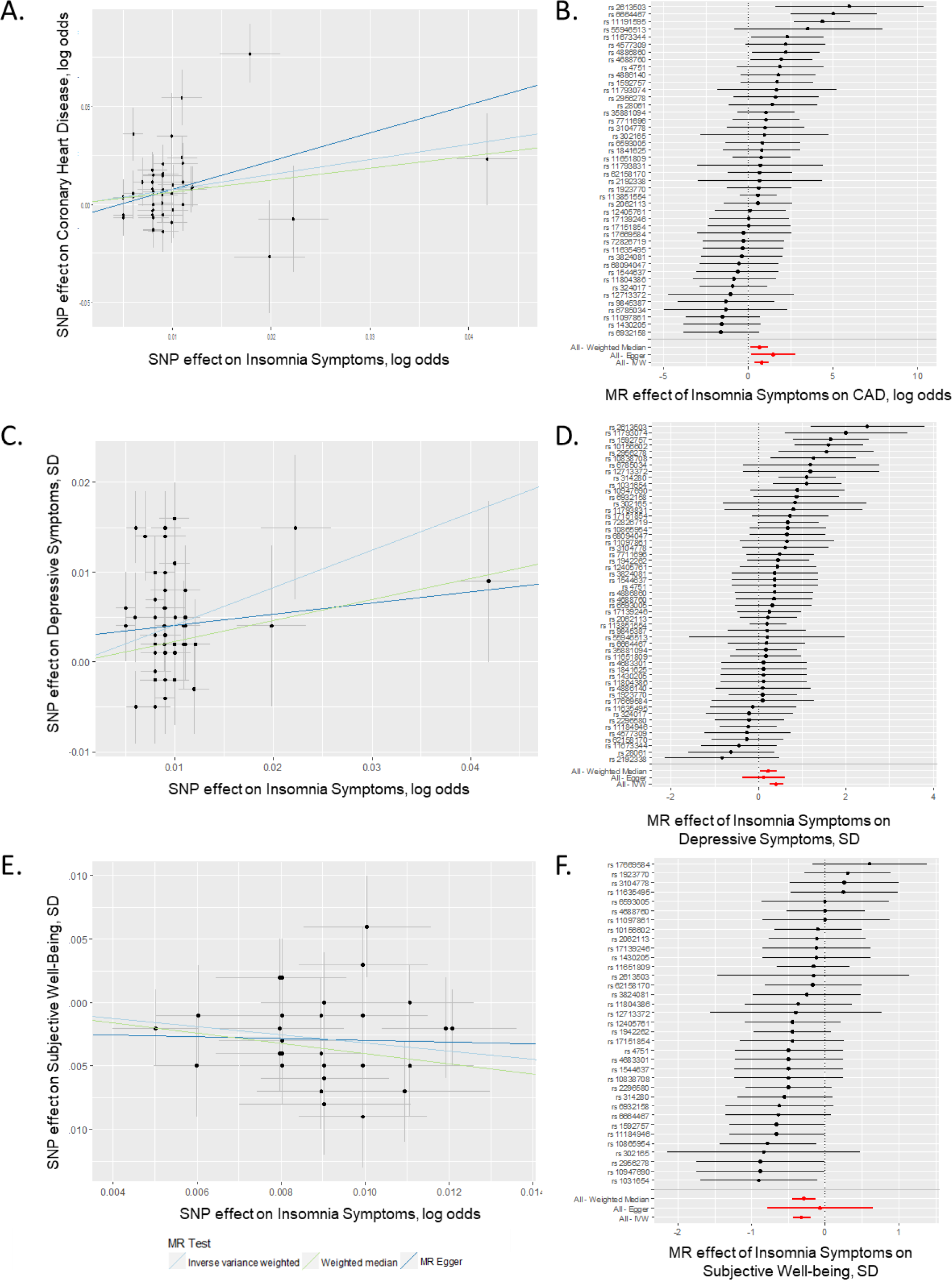
Causal relationships of insomnia symptoms. Association between single nucleotide polymorphisms associated with frequent insomnia symptoms and CAD (A), subjective wellbeing (C), and depressive symptoms (E). Per allele associations with risk plotted against per allele associations with frequent insomnia symptom risk (vertical and horizontal black lines around points show 95% confidence interval for each polymorphism) results are shown for three different MR association tests. Forest plots show the estimate of the effect of genetically increased insomnia risk on CAD (B), Depressive Symptoms (D), and Subjective Well-Being (F) as assessed for each SNP. Also shown for each SNP is the 95% confidence interval (gray line segment) of the estimate and the Inverse Variance MR, MR-Egger, and Weighted Median MR results in red.

This study provides the most comprehensive description to date of the genetic architecture of frequent or persistent insomnia symptoms, pointing to putative causal variants and candidate genes, pathways and tissues for functional studies. Further, we define physiological correlates for insomnia symptoms and meaningful clinical links, including genetic overlap with RLS and a causal link with coronary artery disease.

## Acknowledgements and Funding

This research has been conducted using the UK Biobank Resource. We would like to thank the participants and researchers from the UK Biobank who contributed or collected data. This work was supported by NIH grants R01DK107859 (RS), R21HL121728 (RS), F32DK102323 (JML), R01HL113338 (JML, SR and RS), R01DK102696 (RS and FS), R01DK105072 (RS and FS), T32HL007567(JL), HG003054 (XZ), The University of Manchester (Research Infrastructure Fund), the Wellcome Trust (salary support for DWR and AL) and UK Medical Research Council MC_UU_12013/5 (DAL). The following groups provided summary statistics to LDHub and MR-base: ADIPOGen (Adiponectin genetics consortium), C4D (Coronary Artery Disease Genetics Consortium), CARDIoGRAM (Coronary ARtery DIsease Genome wide Replication and Meta-analysis), CKDGen (Chronic Kidney Disease Genetics consortium), dbGAP (database of Genotypes and Phenotypes), DIAGRAM (DIAbetes Genetics Replication And Meta-analysis), ENIGMA (Enhancing Neuro Imaging Genetics through Meta Analysis), EAGLE (EArly Genetics & Lifecourse Epidemiology Eczema Consortium, excluding 23andMe), EGG (Early Growth Genetics Consortium), GABRIEL (A Multidisciplinary Study to Identify the Genetic and Environmental Causes of Asthma in the European Community), GCAN (Genetic Consortium for Anorexia Nervosa), GEFOS (GEnetic Factors for OSteoporosis Consortium), GIANT (Genetic Investigation of ANthropometric Traits), GIS (Genetics of Iron Status consortium), GLGC (Global Lipids Genetics Consortium), GPC (Genetics of Personality Consortium), GUGC (Global Urate and Gout consortium), HaemGen (haemotological and platelet traits genetics consortium), HRgene (Heart Rate consortium), IIBDGC (International Inflammatory Bowel Disease Genetics Consortium), ILCCO (International Lung Cancer Consortium), IMSGC (International Multiple Sclerosis Genetic Consortium), MAGIC (Meta-Analyses of Glucose and Insulin-related traits Consortium), MESA (Multi-Ethnic Study of Atherosclerosis), PGC (Psychiatric Genomics Consortium), Project MinE consortium, ReproGen (Reproductive Genetics Consortium), SSGAC (Social Science Genetics Association Consortium) and TAG (Tobacco and Genetics Consortium), TRICL (Transdisciplinary Research in Cancer of the Lung consortium), UK Biobank. The Nord-Trøndelag Health Study (The HUNT Study) is a collaboration between HUNT Research Centre (Faculty of Medicine, NTNU, Norwegian University of Science and Technology), Nord-Trøndelag County Council, Central Norway Health Authority, and the Norwegian Institute of Public Health. We are grateful for the contributions from Anne Heidi Skogholt, He Zhang and Hyun Min Kang. We would also like to acknowledge the support given to us by the Genotyping core and Jin Chen. The K.G. Jebesen center for genetic epidemiology is financed by Stiftelsen Kristian Gerhard Jebsen, Faculty of Medicine and Health Sciences Norwegian University of Science and Technology (NTNU) and Central Norway Regional Health Authority. Ben Michael Brumpton and Linn Strand received research grants from The Liaison Committee for education, research and innovation in Central Norway.

## Author Contributions

The study was designed by JML, SJ, ARW, HSD, VVH, KA, HW, SGA, ASL, DWR, TMF, MNW, MKR, and RS. JML, SJ, ARW, HSD, VVH, RB, JT, KA, HW, YS, KP, SMP, JW, TMF, DL, MKR, MNW, and RS participated in acquisition, analysis and/or interpretation of data. JML, HSD, HW and RS wrote the manuscript and all co-authors reviewed and edited the manuscript, before approving its submission. RS is the guarantor of this work and, as such, had full access to all the data in the study and takes responsibility for the integrity of the data and the accuracy of the data analysis.

JW is a consultant for Merck and Flex Pharma. He receives royalties from UpToDate. He has received speaker fees and travel support from Otsuka. He has received research grants from UCB Pharma, NeuroMetrix, NIMH, the RLS Foundation, and Luitpold Pharma. MR has acted as a consultant for GSK, Novo Nordisk, Roche and MSD, and also participated in advisory board meetings on their behalf. MR has received lecture fees from MSD and grant support from Novo Nordisk, MSD and GSK.

## Materials and Methods

### UK Biobank

#### Population and study design

Study participants were from the UK Biobank study, described in detail elsewhere^41^. In brief, the UK Biobank is a prospective study of >500,000 people living in the United Kingdom. All people in the National Health Service registry aged 40-69 and living <25 miles from a study center were invited to participate between 2006-2010. In total 503,325 participants (5%) were recruited from over 9.2 million mailed invitations. Self-reported baseline data was collected by questionnaire and anthropometric assessments were performed. For the current analysis, individuals of non-white ethnicity (n=48,667) were excluded to avoid confounding effects.

#### Insomnia and covariate measures

Study subjects self-reported insomnia symptoms, depression, medication use, age, and sex on a touch-screen questionnaire at baseline assessment. Height and weight were also assessed at baseline. To assess insomnia symptoms, subjects were asked, “Do you have trouble falling asleep at night or do you wake up in the middle of the night?” with responses “never/rarely”, “sometimes”, “usually”, “prefer not to answer”. Subjects who responded “Prefer not to answer” (n=637) were set to missing. We undertook two GWAS, one in which insomnia symptoms were dichotomized into controls (“never/rarely”) and cases with any symptoms (“sometimes” and “usually”); and the second in which participants were dichotomized into controls (“never/rarely”) or frequent insomnia symptoms (“usually”), with those reporting “sometimes” excluded. Additional covariates used in sensitivity analyses included BMI, self-reported and prevalent sleep apnea diagnosis, area deprivation index, alcohol intake, snoring, nap behavior, smoking status, menopause status, weekly physical activity, tea and coffee intake, depression, and extreme stress. BMI was calculated from measured height and weight at baseline visit. Prevalent sleep apnea cases were defined based on primary or secondary ICD10 diagnosis code (G47.3; 391 cases) at the time of baseline assessment. Social deprivation for participant area of residence at the time of recruitment was represented by the Townsend deprivation index, based on national census data immediately preceding participation in the UK Biobank. The Townsend deprivation index was log transformed for use in the analyses. Alcohol intake was self-reported in response to the question “About how often do you drink alcohol?” with answers ranging from “daily or almost daily” to “never”. Snoring was reported in answer to the question “Does your partner or a close relative or friend complain about your snoring?”. Nap behavior was self-reported in response to the question “Do you have a nap during the day” with answers “never/rarely”, “sometimes”, or “usually”. Smoking status was self-reported as past smoking behavior and current smoking behavior, and classified into “current”, “past”, or “never” smoked. Menopause status was reported in answer to the question “Have you had your menopause (periods stopped)?” with answers “yes”, “no”, “not sure - had a hysterectomy”, “not sure – other reason”. Weekly physical activity in total metabolic equivalents per week, total MET-h/week, was calculated using self-reported estimates of type and duration of physical activity. Tea and coffee intake was estimated using the questions “How many cups of tea do you drink each day? (Include black and green tea)” and “How many cups of coffee do you drink each day? (Include decaffeinated coffee),” respectively, with responses in cups/day. Employment status was self-reported in response to the following question “Which of the following describes your current situation?” with responses “in paid employment or self-employed”, “retired”, “looking after home and/or family”, “unable to work because of sickness or disability”, “unemployed”, doing unpair or voluntary work”, “full or part-time student”, “none of the above”, “prefer not to answer”. Marital status was derived from self-reported household occupancy and relatedness data as follows: married/partner was derived from those reporting husband/wife/partner in household with >1 person reported to live in household. Depression was reported in answer to the question “How often did you feel down, depressed or hopeless mood in last 2 weeks?” (cases, n=4,242 based on answers “more than half the days”, or “nearly every day”). Extreme stress was accessed with the question “In the last two years have you experiences any of the following (you can select more than one answer” with responses “serious illness, injury or assault to yourself”, “serious illness, injury or assault of a close relative”, “death of a close relative”, “death of a spouse or partner”, “marital separation/divorce”, “financial difficulties”, “none of the above”. A secondary GWAS further excluded shift workers, sleep and psychiatric medication users, and subjects with chronic and psychiatric illness (n=76,470). Cases of psychiatric disorder was determined using any of the following definitions (derived from Howard et al.): 1) ICD-10 codes for major depressive disorder (F32, F33), bipolar disorder (F30, F31), schizophrenia (F20-F29), autism (F84.0, F84.3, F84.5), intellectual disability (F70.0, F70.1, F70.9), anxiety disorder (F40 - F43), multiple personality disorder (F44.8), or mood disorder (F30 - F39). 2) Self-reported antidepressant, antipsychotic, or anxiolytic use at nurse-led interview.3) Self-reported depression, major depressive disorder, bipolar disorder, or schizophrenia at nurse-led interview at baseline visit. 4) “Broad depression”: responded yes to the question “Have you ever seen a general practitioner (GP) for nerves, anxiety, tension or depression?” and yes to either, “have you ever had a time when you were feeling depressed or down for at least a whole week,” or, “Have you ever had a time when you were uninterested in things or unable to enjoy the things you used to for at least a whole week?” lasting for more than 1 week. 5) Questionnaire-assessed bipolar disorder (Smith algorithm) responded yes to the question, “Have you ever had a period of time lasting at least two days when you were feeling so good, "high", excited or "hyper" that other people thought you were not your normal self or you were so "hyper" that you got into trouble?” or “Have you ever had a period of time lasting at least two days when you were so irritable that you found yourself shouting at people or starting fights or arguments?” for a duration of at least one week and with at least 3 manic/hyper Symptoms^42^. We used the following definitions of medication use- Sleep medications: oxazepam, meprobamate, medazepam, bromazepam, lorazepam, clobazam, chlormezanone, temazepam, nitrazepam, lormetazepam, diazepam, zopiclone, triclofos, methyprylone, prazepam, triazolam, ketazolam, dichloralphenazone, clomethiazole, zaleplon, butobarbital. Antidepressants: amitriptyline, citalopram, fluoxetine, sertraline, venlafaxine, dosulepin, paroxetine, mirtazapine, escitalopram, trazodone, prozac, seroxat, cipralex, duloxetine, lofepramine, clomipramine, nortriptyline, imipramine, dothiepin, cipramil, amitriptyline, prothiaden, trimipramine, lustral, reboxetine, zispin, cymbalta, anafranil, doxepin, moclobemide, phenelzine, fluvoxamine, yentreve, triptafen, surmontil, tranylcypromine, allegron, edronax, molipaxin, mianserin, nardil, faverin, nefazodone, amitriptyline+chlordiazepoxide, isocarboxazid, manerix, maoi, sinequan, tranylcypromine+trifluoperazine, ludiomil, norval, tryptizol, and fluphenazine hydrochloride+nortriptyline. Antipsychotics: prochlorperazine, olanzapine, quetiapine, risperidone, chlorpromazine, trifluoperazine, amisulpride, sulpiride, seroquel, haloperidol, aripiprazole, stelazine, depixol, flupentixol, clozapine, promazine, risperdal, modecate, fluanxol, flupenthixol, zyprexa, zuclopenthixol, clopixol, largactil, abilify, fluphenazine, haldol, serenace, clozaril, cpz, perphenazine, levomepromazine, pericyazine, dolmatil, fentazin, fluphenazine, benperidol, pimozide, zaponex, denzapine, neulactil, thioridazine, dozic, fluspirilene, panadeine, and sertindole. Anxiolytics: zopiclone, diazepam, temazepam, zolpidem, nitrazepam, lorazepam, hydroxyzine, zimovane, phenergan, promethazine, buspirone, atarax, oxazepam, loprazolam, chlordiazepoxide, lormetazepam, ucerax, stilnoct, diazepam, buspar, alprazolam, librium, xanax, meprate, dalmane, clomethiazole, meprobamate, welldorm, amitriptyline+chlordiazepoxide, flurazepam, heminevrin, medazepam, neulactil, sinequan, almazine, atensine, carisoma, chloractil, chloral, dichloralphenazone, dormonoct, methyprylone, mogadon, rohypnol and tryptizol.

#### Activity-monitor derived measures of sleep

Actigraphy devices (Axivity AX3) were worn 2.8 - 9.7 years after study baseline by 103,711 individuals from the UK Biobank for up to 7 days. Details are described elsewhere [REF: PMID: 28146576]. Of these 103,711 individuals, we excluded 11,067 individuals based on accelerometer data quality. Samples were excluded if they satisfied at least one of the following conditions (see also http://biobank.ctsu.ox.ac.uk/crystal/label.cgi?id=1008): a non-zero or missing value in data field 90002 (“Data problem indicator”), “good wear time” flag (field 90015) set to 0 (No), “good calibration” flag (field 90016) set to 0 (No), “calibrated on own data” flag (field 90017) set to 0 (No) or overall wear duration (field 90051) less than 5 days. Additionally, samples with extreme values of mean sleep duration (<3 hours or >12 hours) or mean number of sleep periods (<5 or >30) were excluded. 85,502 samples remained after nonwhite ethnicity exclusions. Sleep measures were derived by processing raw accelerometer data (.cwa). First we converted .cwa files available from the UK Biobank to .wav files using Omconvert (https://github.com/digitalinteraction/openmovement/tree/master/Software/AX3/omconvert) for signal calibration to gravitational acceleration^43,44^ and interpolation^44^ The .wav files were processed with the R package GGIR to infer activity monitor wear time^45^, and extract the z- angle across 5-second epoch time-series data for subsequent use in estimating the sleep period time window (SPT-window) [Vincent Van Hees et al. BioRxiv, 2018] and sleep episodes within it^46^.

The SPT-window was estimated using an algorithm described in [Vincent Van Hees et al. BioRxiv, 2018], implemented in the GGIR R package and validated using PSG in an external cohort. Briefly, for each individual, median values of the absolute change in z-angle (representing the dorsal-ventral direction when the wrist is in the anatomical position) across 5- minute rolling windows were calculated across a 24-hour period, chosen to make the algorithm insensitive to activity-monitor orientation. The 10^th^ percentile was incorporated into the threshold distinguishing movement from non-movement. Bouts of inactivity lasting ≥30 minutes are recorded as inactivity bouts. Inactivity bouts that are <60 minutes apart are combined to form inactivity blocks. The start and end of longest block defines the start and end of the SPT-window [van Hees et al 2018, in press]. *Sleep duration.* Sleep episodes within the SPT-window were defined as periods of at least 5 minutes with no change larger than 5° associated with the z-axis of the accelerometer, as motivated and described in van Hees et al.^46^. The summed duration of all sleep episodes was used as indicator of sleep duration. *Sleep efficiency.* This was calculated as sleep duration (defined above) divided by the time elapsed between the start of the first inactivity bout and the end of the last inactivity bout (which equals the SPT-window duration). *Number of sleep episodes within the SPT-window.* This is defined as the number of sleep episodes separated by last least 5 minutes of wakefulness within the SPT-window. The least-active five hours (L5) and the most-active ten hours *(M10)* of each day were defined using a five-hour and ten-hour daily period of minimum and maximum activity, respectively. These periods were estimated using a rolling average of the respectively time window. L5 was defined as the number of hours elapsed from the previous midnight whereas M10 was defined as the number of hours elapsed from the previous midday. *Sleep midpoint* was calculated for each sleep period as the midpoint between the start of the first detected sleep episode and the end of the last sleep episode used to define the overall SPT-window (above). This variable is represented as the number of hours from the previous midnight, e.g. 2am = 26. *Diurnal inactivity duration* is the total daily duration of estimated bouts of inactivity that fall outside of the SPT-window. All activity-monitor phenotypes were adjusted for age at accelerometer wear, sex, season of wear, release (categorical; UK BiLeVe, UKB Axiom interim, release UKB Axiom full release) and number of valid recorded nights (or days for M10) when performing the association test in BOLT-LMM.

#### Coronary Artery Disease

Coronary artery disease (CAD) in the UK Biobank was defined as a diagnosis of myocardial infarction or coronary revascularization, as described previously^47^. Myocardial infarction was defined as a self-reported diagnosis of “heart attack” or “myocardial infarction” or hospitalization/death due to an ICD-10 code for acute myocardial infarction (I21.0, I21.1, I21.2, I21.4, I21.9). Coronary revascularization was defined based on a self-report of “coronary artery bypass grafting” or “coronary artery angioplasty,” or hospitalization for an OPCS-4 code for coronary artery bypass grafting (K40.1-K40.4, K41.1-K41.4, K45.1-K45.5) or coronary artery angioplasty ± stenting (K49.1–49.2, K49.8–49.9, K50.2, K75.1–75.4, K75.8–75.9). Individuals with self-reported “angina” or hospitalization for an ICD-10 code of angina pectoris (I20) or chronic ischemic heart disease (I25.1, I25.5, I25.6, I25.8, I25.9) were excluded from analyses. All other individuals were defined as controls.

#### Genotyping and quality control

Phenotype data is available for 502,631 subjects in the UK Biobank. Genotyping was performed by the UK Biobank, and genotyping, quality control, and imputation procedures are described in detail here^48^. In brief, blood, saliva, and urine was collected from participants, and DNA was extracted from the buffy coat samples. Participant DNA was genotyped on two arrays, UK BiLEVE and UKB Axiom with >95% common content and genotypes for ~800,000 autosomal SNPs were imputed to two reference panels. Genotypes were called using Affymetrix Power Tools software. Sample and SNPs for quality control were selected from a set of 489,212 samples across 812,428 unique markers. Sample QC was conducted using 605,876 high quality autosomal markers. Samples were removed for high missingness or heterozygosity (968 samples) and sex chromosome abnormalities (652 samples). Genotypes for 488,377 samples passed sample QC (~99.9% of total samples). Marker based QC measures were tested in the European ancestry subset (n=463,844), which was identified based on principal components of ancestry. SNPs were tested for batch effects (197 SNPs/batch), plate effects (284 SNPs/batch), Hardy-Weinberg equilibrium (572 SNPs/batch), sex effects (45 SNPs/batch), array effects (5417 SNPs), and discordance across control replicates (622 on UK BiLEVE Axiom array and 632 UK Biobank Axiom array) (p-value <10^-12^ or <95% for all tests). For each batch (106 batches total) markers that failed at least one test were set to missing. Before imputation, 805,426 SNPs pass QC in at least one batch (>99% of the array content). Population structure was captured by principal component analysis on the samples using a subset of high quality (missingness <1.5%), high frequency SNPs (>2.5%) (~100,000 SNPs) and identified the sub-sample of white British descent. We further clustered subjects into four ancestry clusters using K-means clustering on the principal components, identifying 453,964 subjects of European ancestry. Imputation of autosomal SNPs was performed to UK10K haplotype, 1000 Genomes Phase 3, and Haplotype Reference Consortium (HRC) with the current analysis using only those SNPs imputed to the HRC reference panel. Autosomal SNPs were pre-phased using SHAPEIT3^49^ and imputed using IMPUTE4. In total ~96 million SNPs were imputed. Related individuals were identified by estimating kinship coefficients for all pairs of samples, using only markers weakly informative of ancestral background. In total there are 107,162 related pairs comprised of 147,731 individuals related to at least one other subject in the UK Biobank.

#### Genome-wide association analysis

Genetic association analysis was performed in related subjects of European ancestry (n=453,964) using BOLT-LMM^24^ linear mixed models and an additive genetic model adjusted for age, sex, 10 PCs, genotyping array and genetic correlation matrix[jl2] with a maximum per SNP missingness of 10% and per sample missingness of 40%. We used a genome-wide significance threshold of 5×10^-8^ for each GWAS. Genetic association analysis was also performed in unrelated subjects of white British ancestry (n=337,545) using PLINK^50^ logistic regression and an additive genetic model adjusted for age, sex, 10 PCs and genotyping array to determine SNP effects on self-reported insomnia symptoms. We used a hard-call genotype threshold of 0.1, SNP imputation quality threshold of 0.80, and a MAF threshold of 0.001. Genetic association analysis for the X chromosome was performed using the genotyped markers on the X chromosome with the additional –sex flag in PLINK. We are 80% powered to detect a relative difference of 4% or more (i.e. an OR of 1.04 or 0.96 assuming a MAF 0.1, p=5×10^-8^). Sex-specific GWAS were performed in PLINK 1.9^86^ using logistic regression stratified by sex adjusting for age, 10 principal components of ancestry, and genotyping array. We used a hard-call genotype threshold of 0.1, SNP imputation quality threshold of 0.80, and a MAF threshold of 0.001. SNP x sex interactions (13 tests) and SNP x menopause interaction in females (13 tests) were tested for genome-wide significant signals with the significance threshold defined by Bonferroni correction. Trait heritability was calculated as the proportion of trait variance due to additive genetic factors measured in this study using BOLT-REML^24^, to leverage the power of raw genotype data together with low frequency variants (MAF≥0.001).

#### Post-GWAS analyses

##### Sensitivity analyses on top signals

Follow-up analyses on genome-wide significant loci in the primary analyses included covariate sensitivity analysis individually adjusting for sleep apnea, coffee/tea intake, physical activity, severe stress, depression, psychiatric medication use, socio-economic status, smoking, employment and marital status, and snoring, or BMI in addition to baseline adjustments for age, sex, 10 PCs and genotyping array. Sensitivity and sex-specific analyses were conducted only in unrelated subjects of white British ancestry.

##### Gene, pathway and tissue-enrichment analyses

Gene-based analysis was performed using PASCAL^20^. Tissue enrichment analysis was conducted using FUMA^51^. Enrichment for pathways and ontologies was performed in EnrichR^27,28^ using the human genome as the reference set and a minimum number of 2 genes per category. A genetic risk score for Restless Legs Syndrome (RLS) was tested using the weighted genetic risk score calculated by summing the products of the RLS risk allele count for 20 genome-wide significant SNPs multiplied by the scaled RLS effect reported by Schormair et al.^34^ using the summary statistics from our frequent insomnia symptom GWAS using the GTX package in R^52^. Integrative transcriptome-wide association analyses with GWAS were performed using the FUSION TWAS package^23^ with weights generated from gene expression in 8 brain regions and 6 tissues from the GTEX consortium (v6). Tissues for TWAS testing were selected from the FUMA tissue enrichment analyses and here we present significant results that survive Bonferroni correction for the number of genes tested per tissue and for all 14 tissues.

##### Genetic correlation analyses

Post-GWAS genome-wide genetic correlation analysis of LD Score Regression (LDSC)^25,35,36^ using LDHub was conducted using all UK Biobank SNPs also found in HapMap3 and included publicly available data from 224 published genome-wide association studies, with a significance threshold of *p*=0.002 after Bonferroni correction for all tests performed. LDSC estimates genetic correlation between two traits from summary statistics (ranging from -1 to 1) using the fact that the GWAS effect-size estimate for each SNP incorporates effects of all SNPs in LD with that SNP, SNPs with high LD have higher X^2^ statistics than SNPs with low LD, and a similar relationship is observed when single study test statistics are replaced with the product of z-scores from two studies of traits with some correlation. Furthermore, genetic correlation is possible between case/control studies and quantitative traits, as well as within these trait types. We performed partitioning of heritability using the 8 pre-computed cell-type regions, and 25 pre-computed functional annotations available through LDSC, which were curated from large-scale robust datasets^25^. Enrichment both in the functional regions and in an expanded region (+500bp) around each functional class was calculated in order to prevent the estimates from being biased upward by enrichment in nearby regions. The multiple testing threshold for the partitioning of heritability was determined using the conservative Bonferroni correction (p of 0.05/25 classes). Summary GWAS statistics will be made available at the UK Biobank web portal.

##### Mendelian randomization analyses

MR analysis was carried out using MR-Base^53^ (ref), using the inverse variance weighted approach as our main analysis method^37^, and MR-Egger^38^ and weighted median estimation^39^ as sensitivity analyses. MR results may be biased by horizontal pleiotropy – i.e. where the genetic variants that are robustly related to the exposure of interest (here frequent insomnia symptoms) independently influence levels of a causal risk factor for the outcome. IVW assumes that there is no horizontal pleiotropy. MR-Egger provides unbiased causal estimates even if all of the genetic instruments have horizontal pleiotropic effects, but it has an additional assumption that the association of genetic instruments with risk factor is not correlated with pleiotropic genetic instrument association with outcome. The weighted median approach is valid if less than 50% of the weight is pleiotropic (i.e. no single SNP that contributes 50% of the weight or a number of SNPs that together contribute 50% should be invalid because of horizontal pleiotropy. Given these different assumptions, if all three methods are broadly consistent this strengthens our causal inference. For most of our MR analyses we used two-sample MR, in which, for all 57 of the insomnia GWAS hits identified in this study, we looked for the per allele difference in odds (binary outcomes) or means (continuous) with outcomes from summary publicly available data in the MR-Base platform. Results are therefore a measure of ‘any insomnia’ and sample 1 is UK Biobank (our GWAS) and sample 2 a number of different GWAS consortia covering the outcomes we explored (Supplementary Table 17). For all four of the longevity outcomes and as follow up for CAD we used one-sample MR with the SNP-outcome associations also obtained from UK Biobank. If we could not find one of the 57 SNPs in the outcome database we substituted for a proxy where possible, LD proxies are defined using 1000 genomes European sample with R^2^ >0.8. The number of SNPs used in each MR analysis varies by outcome from 11 to 53 because of some SNPs (or proxies for them) not being located in the outcome GWAS (Table 1).

Primary association analyses of the 57 genome-wide significant Insomnia SNPs with CAD were performed in Hail (https://github.com/hail-is/hail) using imputed genotype dosages and a logistic regression model adjusting for age at first visit, sex, genotyping array, and the first 10 principal components of ancestry. A total of 23,980 CAD cases were compared to 388,326 referents. For Mendelian Randomization, a fixed-effects inverse-variance weighted meta-analysis was performed of the SNP-specific association estimates with CAD, aligning each Insomnia SNP allele/beta coefficient to “increased risk of insomnia.” Sensitivity analyses were performed excluding the variants with the strongest effect estimate and/or widest confidence intervals to account for SNP heterogeneity.

### Replication Cohorts

#### The HUNT Study

##### Sample ascertainment and phenotype definition

The Nord-Trøndelag Health Study (HUNT) consists of three different population-based health surveys conducted in the county of Nord-Trøndelag, Norway over approximately 20 years (HUNT1 [1984-1986], HUNT2 [1995-1997] and HUNT3 [2006-2008])^14^. At each survey, the entire adult population (≥ 20 years) was invited to participate by completing questionnaires, attending clinical examinations and interviews. Participation rates in HUNT1, HUNT2 and HUNT3 were 89.4% (n=77,212), 69.5% (n=65 237) and 54.1% (n=50 807), respectively^14^. Taken together, the study included more than 120,000 different individuals from Nord-Tr0ndelag County. Biological samples including DNA have been collected for approximately 70,000 participants. The HUNT Study has been described in more detail elsewhere^14^

Participants reporting insomnia in either HUNT1, HUNT2 or HUNT3 were classified as having insomnia for the present study. In HUNT1, participants who answered “often” or “almost every night” to the question “During the last month, have you had any problems falling asleep or sleep disorders?” were classified as having insomnia. Those who answered “almost every night” were additionally classified as having frequent insomnia. In HUNT2, participants who answered “often” or “almost every night” to either of the questions “Have you had difficulty falling asleep in the last month?” or “During the last month, have you woken too early and not been able to get back to sleep?”, were classified as having insomnia. Those who answered “almost every night” to either question were additionally classified as having frequent insomnia. In HUNT3, participants were asked how often in the last 3 months they had “had difficulty falling asleep at night” and “woken up repeatedly during the night” with the response options “Never/seldom”, “Sometimes” and “Several times a week”. Those who answered “Several times a week” to either question were classified as having insomnia. Because of fewer response options on the insomnia questions in HUNT3 than in HUNT1 and HUNT2, we classified frequent insomnia differently than in the two previous studies, and only those who answered “Several times a week” to both questions were classified as having frequent insomnia.

Participants were classified as controls (no insomnia) if they answered the insomnia questions in at least one of the three HUNT studies, and did not qualify as cases.

##### Genotyping, quality control and imputation

DNA from 71,860 HUNT samples was genotyped using one of three different Illumina HumanCoreExome arrays (HumanCoreExome12 v1.0, HumanCoreExome12 v1.1 and UM HUNT Biobank v1.0). Genotyping and quality control are described here [PMID:29083406]. Imputation was performed on samples of recent European ancestry using Minimac3 (v2.0.1, http://genome.sph.umich.edu/wiki/Minimac3)^54^ and a merged reference panel that was constructed by combining the Haplotype Reference Consortium panel (release version 1.1)^55^ and a local reference panel based on 2,202 whole-genome sequenced HUNT study participants. After restricting to the above, those with European ancestry and with available phenotype information, 62,533 individuals were included in the analysis sample.

##### Association analysis

Association analyses were conducted using SAIGE [Efficiently controlling for case-control imbalance and sample relatedness in large-scale genetic association studies. bioRxiv, doi: https://doi.org/10.1101/212357], a generalized mixed effects model approach, to account for cryptic population structure and relatedness when modelling the association between genotype probabilities (dosages) and insomnia symptoms. Models were adjusted for sex, birth year, genotyping batch and four principal components (PCs). PCs were computed using PLINK. Additional filters applied to the analysis included minor allele count ≥ 5 and imputation r2 ≥ 0.3.

##### Partners Healthcare Biobank

The Partners Biobank is an ongoing hospital-based cohort study launched in 2010 from the Partners HealthCare hospitals with electronic medical record (EMR) and genetic data supplemented with health surveys. Recruitment is from participating clinics at Brigham and Women’s Hospital (BWH), Massachusetts General Hospital (MGH), Spaulding Rehabilitation Hospital (SRH), Faulkner Hospital (FH) and McLean Hospital (MCL), (NWH), (NSM) and electronically from previous patients. At the time of analysis, a total of 73,683 participants have provided consent, of which 20,087 have been genotyped. Cases of insomnia were ascertained from EMR using diagnoses of insomnia (and primary insomnia) (n =2,217). Participants without insomnia and restless leg syndrome were selected as controls (n =17,018). Participants were genotyped using the Illumina Multi-Ethnic GWAS/Exome SNP Array. Imputation was performed using Minimac3 (http://genome.sph.umich.edu/wiki/Minimac3) using the HRC (Version r1.1 2016) reference panel for imputation. This HRC panel consists of 64,940 haplotypes of predominantly European ancestry. Haplotype phasing was performed using SHAPEIT^56^. In total, 16,791 samples of self-reported European ancestry with high-quality genotyping, EMR data and covariate data were used for these analyses. We derived a genetic risk score (GRS) for frequent insomnia symptoms using 57 SNPs. Individual participant scores were created by summing the number of risk alleles at each genetic variant, which where weighted by the respective allelic effect sizes on frequent insomnia symptoms. We tested whether the GRSs for insomnia by estimating linear trends of the GRS adjusted for age, sex, and genotyping array.

Summary GWAS statistics will be made available at the UK Biobank website (http://biobank.ctsu.ox.ac.uk/) and through the Saxena lab webpage (https://cgm.massgeneral.org/faculty/richa-saxena/).

## Supplementary Materials

Supplementary Figure. 1-6

Supplementary Table. 1-20

